# An upstream open reading frame represses translation of chicken PPARγ transcript variant 1

**DOI:** 10.1101/858753

**Authors:** Yankai Chu, Jiaxin Huang, Guangwei Ma, Tingting Cui, Xiaohong Yan, Hui Li, Ning Wang

## Abstract

Peroxisome proliferator-activated receptor γ (PPARγ) is a master regulator of adipogenesis. The *PPARγ* gene produces various transcripts with different 5′-untranslated regions (5′ UTRs) because of alternative promoter usage and splicing. The 5′ UTR plays important roles in posttranscriptional gene regulation. However, to date, the regulatory role and underlying mechanism of 5′ UTRs in the posttranscriptional regulation of PPARγ expression remain largely unclear. In this study, we investigated the effects of 5′ UTRs on posttranscriptional regulation using reporter assays. Our results showed that the five PPARγ 5′ UTRs exerted different effects on reporter gene activity. Bioinformatics analysis showed that chicken PPARγ transcript 1 (*PPARγ1*) possessed an upstream open reading frame (uORF) in its 5′ UTR. Mutation analysis showed that a mutation in the uORF led to increased *Renilla* luciferase activity and PPARγ protein expression, but decreased *Renilla* luciferase and *PPARγ1* mRNA expression. mRNA stability analysis using real-time RT-PCR showed that the uORF mutation did not interfere with mRNA stability, but promoter activity analysis of the cloned 5′ UTR showed that the uORF mutation reduced promoter activity. Furthermore, *in vitro* transcription/translation assays demonstrated that the uORF mutation markedly increased the translation of *PPARγ1* mRNA. Collectively, our results indicate that the uORF represses the translation of chicken *PPARγ1* mRNA.

## Introduction

Peroxisome proliferator-activated receptor γ (PPARγ) is a member of the PPAR subfamily of ligand-activated transcription factors. *In vitro* and *in vivo* studies have demonstrated that PPARγ is essential for adipocyte differentiation, adipocyte survival, adipocyte function, insulin sensitivity, and lipogenesis (Lehrke and Lazar 2005; Lefterova *et al.* 2014). Synthetic PPARγ agonists have been used as therapeutic agents for diabetes and insulin insensitivity (Cariou *et al.* 2012).

The *PPARγ* gene is controlled by multiple alternative promoters (Aprile *et al.* 2014; Chandra *et al.* 2017). Because of alternative promoter usage and splicing, the *PPARγ* gene can produce multiple transcript variants, resulting in expression of two PPARγ protein isoforms that differ in the N-terminal. All PPARγ transcript variants differ in their 5′-untranslated regions (5′ UTRs) (Mcclelland *et al.* 2014). These PPARγ 5′ UTR isoforms have distinct tissue distributions (Ahmadian *et al.* 2013), suggesting that the 5′ UTRs may be involved in posttranscriptional and translational regulation of the *PPARγ* gene.

The 5′ UTRs of mRNAs exert crucial roles in posttranscriptional and translational regulation. Several *cis*-regulatory elements within the 5′ UTRs have been identified, such as the 5′ cap structure (Mitchell *et al.* 2010), upstream open reading frames (uORFs) (Hood *et al.* 2009; Barbosa *et al.* 2013), internal ribosome entry sites (IRES) (Xia and Holcik 2009), terminal oligo-pyrimidine tracts, secondary structures, and G-quadruplexes (Yamashita *et al.* 2008; Bugaut and Balasubramanian 2012). These *cis*-regulatory elements can function via various mechanisms, controlling mRNA stability (Nasif *et al.* 2018), nuclear export, localization, and translation efficiency (Araujo *et al.* 2012). Of these *cis*-regulatory elements, uORFs have been widely studied. Bioinformatics analysis showed that about 50% of human transcripts contain uORFs (Suzuki *et al.* 2000; Iacono *et al.* 2005; Calvo *et al.* 2009), and experimental studies have revealed that a number of uORFs can affect the expression of the main downstream ORFs by inducing mRNA decay or by regulating translation (Iacono *et al.* 2005; Crowe *et al.* 2006; Sathirapongsasuti *et al.* 2011).

Given the importance of PPARγ in various physiological and pathological processes, *PPARγ* gene regulation has been extensively studied at the genomic and transcriptional levels in recent decades (Lee and Ge 2014). The half-life of PPARγ mRNA and protein is short and PPARγ protein can be posttranslationally modified in various ways (van Beekum *et al.* 2009; Katsura *et al.* 2014), suggesting that posttranscriptional regulation is crucial for its function. However, to date, posttranscriptional regulation by the 5′ UTR has been mostly unexplored. In the present study, we investigated the posttranscriptional regulation of chicken PPARγ by the 5′ UTR. Of interest, we demonstrated that translation of chicken PPARγ transcript variant 1 (*PPARγ1*) is repressed by a uORF that is absent in human and mouse PPARγ transcripts.

## Materials and Methods

### Cell culture

DF1 cells were purchased from the Institute of Biochemistry and Cell Biology, Chinese Academy of Sciences, and the immortalized chicken preadipocyte cell line 1 (ICP1) was generated in our laboratory (Wang *et al.* 2017). All cells were grown in Dulbecco’s modified Eagle’s medium supplemented with 10% fetal bovine serum and 1% penicillin/streptomycin, at 37 °C and 5% CO_2_. The culture medium was changed two to three times per week and cells were passaged 1:3 or 1:5 as needed.

### Plasmid construction

The reporter vector psi-CHECK2 (Invitrogen, Carlsbad, CA, USA) was modified by site-directed mutagenesis, such that the ATG of *Renilla* luciferase was mutated to TTG, and the resulting reporter vector was named psi-CHECK2-Mut. The DNA sequences corresponding to the five PPARγ 5′ UTRs plus initiation codon ATG were synthesized and inserted into the *Nhe*I restriction site upstream of the *Renilla* luciferase gene. Thus, *Renilla* luciferase in psi-CHECK2-Mut would be expressed with PPARγ with indicated 5′ UTRs and translated with the initiation codon ATG of PPARγ.

To test the promoter activity of the DNA sequence corresponding to chicken PPARγ1 5′ UTR, the wild-type or uORF-mutant 5′ UTRs were subcloned into the *Bam*HI and *Xho*I restriction sites of pGL3-basic vector and named pGL3-PPARγ-WT and pGL3-PPARγ-Mut, respectively.

For PPARγ expression constructs, the full-length coding sequence of PPARγ1 was PCR amplified from the cDNA derived from DF-1 cells with a set of primers (forward primer: 5′-GAATTCATGGTTGACACAGAAATGCCGT-3′ and reverse primer: 5′-CCTCGAGGAGGATAAGAACTACTATCGCC-3′ and cloned into the *Bam*HI and *Eco*RI restriction sites of the pcDNA3.1 expression vector. The synthesized wild-type or uORF mutated 5′ UTR of PPARγ was inserted upstream of PPARγ ORF in pcDNA3.1 vector with *Nhe*I restriction sites and named pcDNA3.1-PPARγ-WT and pcDNA3.1-PPARγ-Mut, respectively. All constructs were confirmed by DNA sequencing and restriction enzyme digestion.

### Quantitative real-time PCR assays (qRT-PCRs)

Total RNA was isolated from ICP1 or DF1 cells by using the RNeasy Plus Mini Kit (Qiagen, Hilden, Germany) according to the manufacturer’s protocol, and the first-strand cDNA was synthesized from 1 μg of total RNA with oligo dT or random primers using ImProm-II reverse transcriptase (Promega, Madison, WI, USA). The qPCR reactions were performed in a 20 μL reaction mixture using SYBR Green PCR Master Mix (Roche, Madison, WI, USA). The primers were as follows: *hRluc* (forward 5′-TGATCGAGTCCTGGGACGA-3′, reverse 5′-ACAATCTGGACGACGTCGGG-3′); wild-type and uORF-mutated PPARγ1 (forward 5′-GGAGTTTATCCCACCAGAAG-3′, reverse 5′-AATCAACAGTGGTAAATGGC-3′); *NONO* (forward 5′-AGAAGCAGCAGCAAGAAC-3′, reverse 5′-TCCTCCATCCTCCTCAGT-3′). qPCR was carried out in an ABI 7500 real-time PCR system (Applied Biosystems, Foster City, CA, USA), and PCR results were recorded as threshold cycle numbers (Ct). The fold change in the target gene expression, normalized to the expression of an internal control gene (*NONO*) and relative to the expression at time point 0 (Normann *et al.* 2016), was calculated using the 2^−ΔΔCT^ method (Livak and Schmittgen 2001). The results are presented as the mean ± SEM of three independent experiments.

### Protein isolation and western blot analysis

The ICP1 cells were transfected with either pcDNA3.1-PPARγ-WT or pcDNA-PPARγ-Mut vector. At 48 h post-transfection, cells were washed twice with PBS and lysed using radioimmunoprecipitation assay (RIPA) buffer (Beyotime Institute of Biotechnology, Beijing, China) supplemented with 1% protease inhibitor mixture. Equal amounts of protein extracts were separated by sodium dodecyl sulfate-PAGE, transferred onto Immun-Blot PVDF membranes (Millipore, Billerica, MA, USA). The membrane was blocked for 1 to 2 h at room temperature with Tris-buffered saline containing 0.1% Tween and 5% non-fat dry milk, and immunoblotted with rabbit polyclonal antibody to chicken PPARγ (1:1000 dilution) or β-actin (1:1000 dilution, ZSGB-BIO, Beijing, China) at room temperature for 1 h. Horseradish peroxidase-conjugated goat anti-mouse or goat anti-rabbit IgG (Promega; 1:10,000) was incubated for 1 h at room temperature and then washed four times with PBS-Tween for 20 min. The immunoreactive bands were visualized using an ECL Plus detection kit (HaiGene Biotechnology, Harbin, China). Immunoreactive protein levels were determined semi-quantitatively by densitometric analysis using the UVP system Labworks TM software 3.0 (UVP, Upland, CA, USA). Each western blot analysis was performed at least three times.

### Dual-luciferase reporter assays

Transient transfections were performed using Lipofectamine 2000 (Invitrogen, Carlsbad, CA, USA). Cells were plated at 1.0 to 1.5 × 10^5^ cells per well in 24-well plates. For the 5′ UTR reporter gene assay, 1 μg of the indicated reporter constructs was transfected into each well. For the promoter reporter gene assay, 0.8 μg of the indicated reporter constructs and 0.4 μg of pRL-TK (Promega), as an internal control of transfection efficiency, were co-transfected into each well. Luciferase activity was analyzed at 48 h post-transfection, using a dual-luciferase reporter kit (Promega) as per the manufacturer’s instructions. All luciferase reporter assays were performed at least three times in quadruplicates.

### *In vitro* transcription and translation

Plasmids pcDNA3.1-PPARγ-WT and pcDNA3.1-PPARγ-Mut were linearized, purified by agarose gel electrophoresis, eluted with diethylpyrocarbonate-treated H_2_O, and quantified. Equal amounts (1 μg) of linearized DNA were used as templates for *in vitro* transcription in the T7 RiboMAX Large Scale RNA Production System (Promega) according to the manufacturer’s protocol. Capped mRNAs were generated using the Ribo m^7^G Cap Analog (Promega). The capped mRNAs were digested with DNase I and purified with the RNeasy kit (Qiagen) and quantified. The size and integrity of the purified mRNAs were assessed by gel electrophoresis. The mRNA outputs of pcDNA3.1-PPARγ-WT and pcDNA3.1-PPARγ-Mut were analyzed by absolute qRT-PCR. *In vitro* translation reactions were performed in nuclease-treated Rabbit Reticulocyte Lysate (Promega) as described by the manufacturer. Equal amounts of the capped mRNA (2 μg) derived from pcDNA3.1-PPARγ-WT or pcDNA3.1-PPARγ-Mut construct were used as the template for *in vitro* translation, which was performed for 60 min at 30 °C, and the reactions were stopped by transferring the tubes to ice. Biotinylated lysine residues were added to the translation reaction as a precharged ε-labeled biotinylated lysine-tRNA complex (Transcend tRNA; Promega) and incorporated into nascent proteins during translation. The translated protein was analyzed using a Transcend Non-Radioactive Translation Detection System (Promega).

### RNA stability assay

The stability of luciferase mRNA transcripts from the indicated constructs (PPARγ1-5′UTR-WT and PPARγ1-5′UTR-Mut) was determined by measuring the amount of *hRluc* luciferase mRNA at selected intervals: 0 (control), 3, 6, 9, and 12 h, following the addition of 5 mg/mL actinomycin D (Sigma-Aldrich, St. Louis, MO, USA) at 48 h post-transfection. Time-course intervals were chosen based on the manufacturer’s data of *luc2* mRNA half-life (approximately 2 h). For mRNA expression analysis, total RNA (1 μg) was reverse-transcribed into cDNA using the PrimeScript RT reagent Kit with gDNA Eraser (Takara, Shiga, Japan), and relative mRNA expression was determined by real-time PCR using FastStart Universal SYBR Green Master [Rox] (Roche) with *hRluc* primers as described above. Relative mRNA levels were normalized to the *NONO* gene and to expression at time point 0 (Normann *et al.* 2016) and calculated using the 2^−ΔΔCT^ method (Livak and Schmittgen 2001).

### Bioinformatics analysis

Online software programs used to predict the potential *cis*-regulatory elements of PPARγ 5′ UTR: StarORF (http://star.mit.edu/index.html) (Ceraj *et al.* 2009), UTRscan (http://itbtools.ba.itb.cnr.it/utrscan) (Grillo *et al.* 2010), and Reg RNA2.0 (http://regrna2.mbc.nctu.edu.tw) (Chang *et al.* 2013). Preliminary RNA secondary structures were predicted using Vienna RNAfold 2.0 (http://rna.tbi.univie.ac.at/cgi-bin/RNAfold.cgi) (Hofacker 2003). Intrinsic protein disorder analyses were made using PSIPRED protein sequence analysis workbench (http://bioinf.cs.ucl.ac.uk/psipred/) (Buchan *et al.* 2013). All bioinformatic computations were performed using default prediction parameters.

### Statistical analysis

Experimental data were analyzed using GraphPad Prism software (GraphPad Inc., San Diego, CA, USA). The results were presented as mean ± SEM. For comparison of two groups, statistical analysis was performed using two-tailed Student’s *t*-test and linear regression. *P* values < 0.05 (*) were considered significant, *P* values < 0.01 (**) were considered highly significant. For multiple comparisons, one-way analysis of variance (ANOVA) was used to determine significance, followed by Tukey’s post hoc test.

### Data availability

Strains, plasmids and cell lines are available upon request. The authors affirm that all data necessary for confirming the conclusions of the article are present within the article, figures and supplemental information.

## Results

### The effects of PPARγ 5′ UTRs on reporter gene expression

We previously identified five different chicken PPARγ transcript variants (*PPARγ* 1 to 5) by 5′ rapid amplification of cDNA ends (5′ RACE) in chicken abdominal adipose tissue (Duan *et al.* 2015). These chicken PPARγ transcript variants encode two protein isoforms (PPARγ1 and PPARγ2) that differ in their N-terminal extension. Chicken PPARγ2 contains 6 additional amino acids at the N-terminus compared with PPARγ1. These five chicken PPARγ transcript variants differed in 5′ UTR sequence and length, and had different tissue distribution patterns (Duan *et al.* 2015), suggesting that 5′ UTRs may play a role in the posttranscriptional regulation of *PPARγ* gene expression. To investigate the posttranscriptional regulatory roles of the five 5′ UTR isoforms on chicken *PPARγ* gene expression, we constructed their respective 5′ UTR reporter constructs. Briefly, we first generated the 5′ UTR reporter vector by mutating the ATG start codon of the *Renilla* luciferase gene to TTG in the psi-CHECK2 vector (Invitrogen) by site-directed mutagenesis (Kubokawa *et al.* 2010); the resulting 5′ UTR reporter construct, named psi-CHECK2-Mut, was used to construct the five chicken PPARγ 5′ UTR reporters. Then, the five DNA fragments corresponding to the five different PPARγ 5′ UTRs plus the ATG start codon were synthesized and inserted at the *Nhe*I restriction site upstream of the *Renilla* luciferase gene in psi-CHECK2-Mut to yield five chicken PPARγ 5′ UTR reporter constructs: PPARγ1-5′UTR, PPARγ2-5′UTR, PPARγ3-5′UTR, PPARγ4-5′UTR, and PPARγ5-5′UTR.

We transfected these five 5′ UTR reporters into ICP1 and DF1 cells and measured *Renilla* luciferase activity. The reporter gene assay showed that these five 5′ UTR reporters displayed different luciferase activities. As shown in Fig. 1A and 1B, PPARγ1-5′UTR exhibited the highest luciferase activity in ICP1 cells and the second-highest activity in DF1 cells. PPARγ5-5′UTR exhibited the highest activity in DF1 cells and the second-highest activity in ICP1 cells. PPARγ3-5′UTR exhibited the lowest activity in both ICP1 and DF1 cells. PPARγ2-5′UTR and PPARγ4-5′UTR exhibited similar reporter activity in both ICP1 and DF1 cells. These results support our speculation that the 5′ UTR regulates chicken *PPARγ* gene expression.

**Figure 1.**
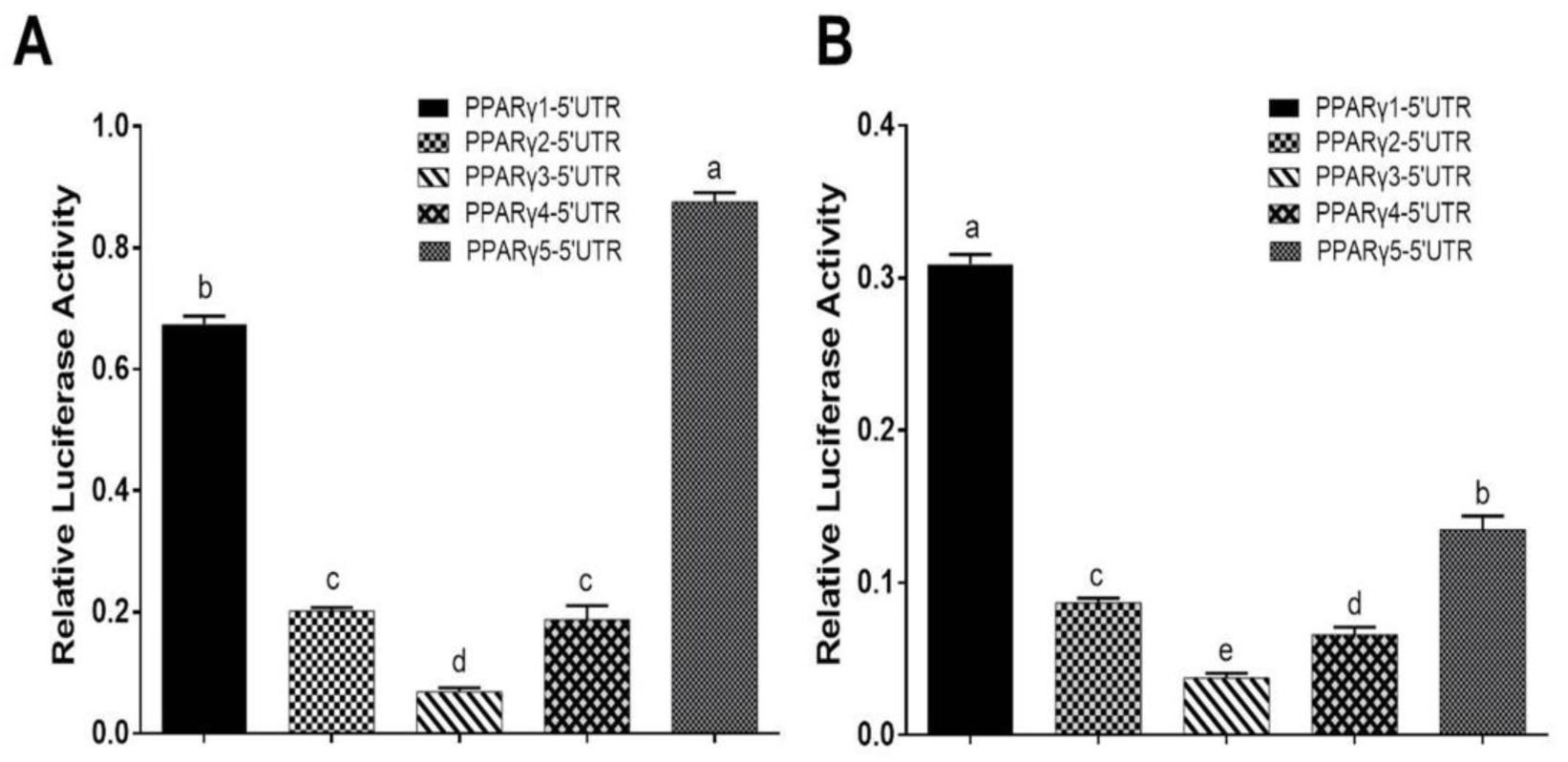
Effects of PPARγ 5′ UTR isoforms on reporter gene activity. (A) The luciferase activity of each of the PPARγ 5′ UTR reporter constructs was measured in DF1 cells. (B) The luciferase activity of each of the PPARγ 5′ UTR reporter constructs was measured in ICP1 cells. Data are expressed as mean ± SEM (n ≥ 3 independent experiments). Bars with different superscripts are statistically different (*P* < 0.05).

### Bioinformatics analysis of chicken PPARγ 5′ UTRs

To gain insight into the molecular mechanisms by which the 5′ UTRs regulate gene expression, we performed a bioinformatics analysis of these five 5′ UTR sequences using the online software programs StarORF, UTRscan, and RegRNA 2.0. Bioinformatics analysis showed that PPARγ1 5′ UTR contains a 54-nucleotide (nt)-long uORF (PPARγ1 uORF), PPARγ3 5′ UTR has a 12-nt-long uORF (PPARγ3 uORF) and a putative IRES element, and PPARγ5 5′ UTR has two uORFs, which are 15 and 51 nt long, respectively. No putative *cis*-regulatory elements were predicted in PPARγ2 and PPARγ4 5′ UTR sequences. The PPARγ1 uORF is located in its 5′ UTR from nucleotides −24 to −79 (relative to the start codon AUG of the PPARγ protein-coding ORF, where A is +1; Fig. 2A), and the uORF AUG (uAUG) resides in a favorable Kozak consensus context, suggesting that there is a high probability that scanning ribosomes consistently initiate the translation at this uAUG codon to encode a 17-amino acid peptide (MGRPGEFIPPEGNSFSG; Fig. 2A).

**Figure 2.**
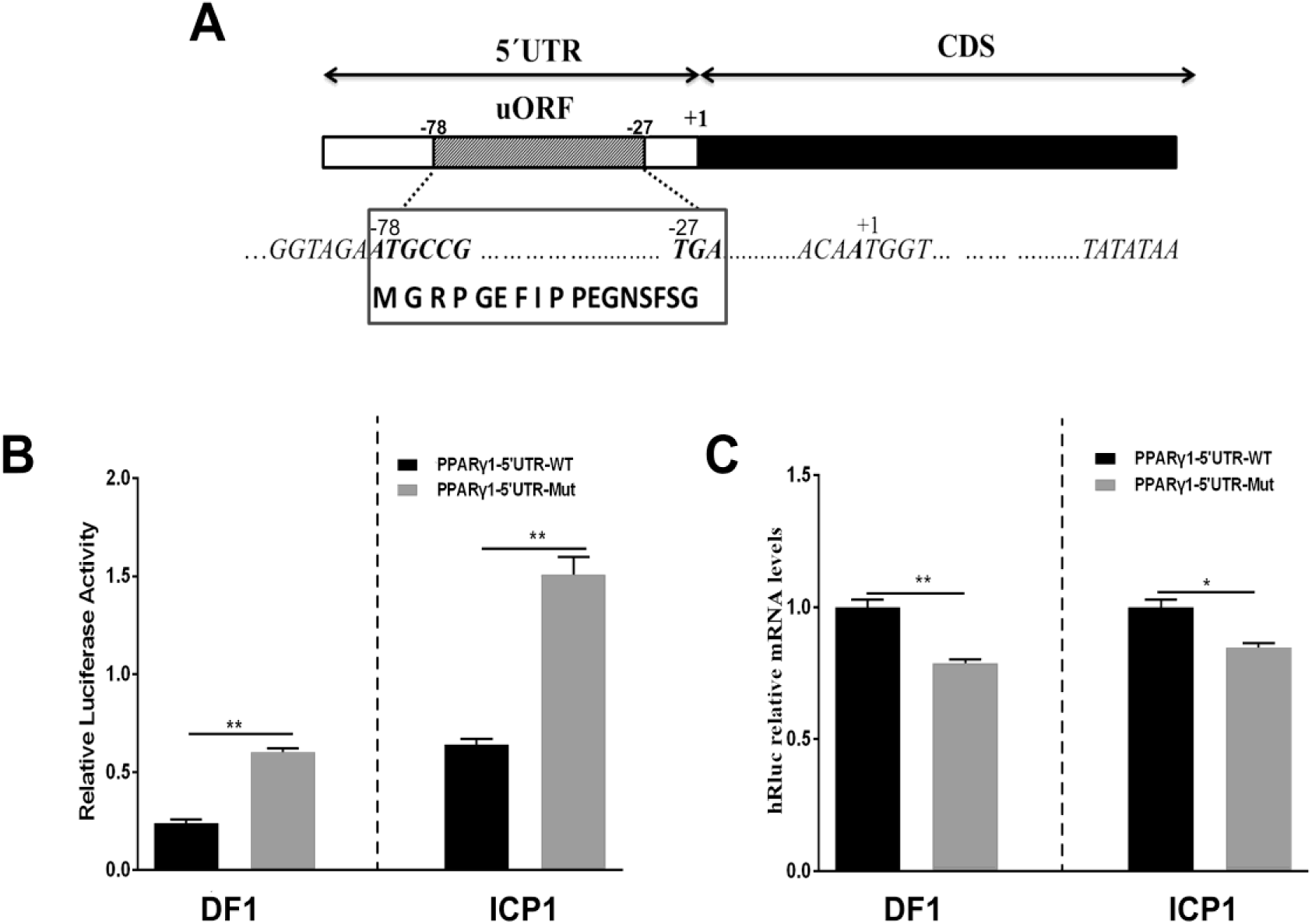
Schematic representation of PPARγ1 5′ UTR and effects of the PPARγ1 uORF mutation on *hRluc* luciferase activity and mRNA expression. (A) A schematic diagram of the 117- nucleotide-long PPARγ1 5′ UTR, the uORF is from nucleotides −24 to −79 of the 5′ UTR, and indicated by a striped rectangle. All positions are numbered relative to the initiation codon ATG of PPARγ transcript 1 (PPARγ1). The uORF encodes a 17-amino acid peptide with the amino acid sequence shown in the bottom. (B) The effect of uORF mutation on the luciferase reporter gene activity. The wild-type (PPARγ1-5′UTR-WT) and uORF mutant (PPARγ1-5′UTR-Mut) PPARγ1 5′ UTR reporter constructs were transfected into ICP1s and DF1 cells, respectively, and reporter gene activity was measured. Compared with the wild-type PPARγ1 5′ UTR reporter, the luciferase activity of PPARγ1-5′UTR-Mut was significantly higher than that of PPARγ1-5′UTR-WT in both ICP1 and DF1 cells (n ≥ 3, ** *P* < 0.01, Student’s *t*-test). (C) The *hRluc* mRNA quantification by real-time RT-PCR in the ICP1 and DF1 cells transfected with the indicated reporter constructs. The relative *hRluc* mRNA Levels are normalized to the expression levels of the cells transfected with the reporter PPARγ1-5′UTR-WT. Data were expressed as the mean ± SEM, *NONO* was used as the internal mRNA control. N ≥ 3, \**P* < 0.05; ** *P* < 0.01, Student’s *t*-test.

### The effect of PPARγ1 uORF on gene expression

Upstream ORFs have emerged as a major posttranscriptional regulatory element in eukaryotic species (Wen *et al.* 2009). The above bioinformatics analysis showed that, of these five PPARγ 5’UTR isoforms, three contained uORFs, which led us to speculate that these uORFs may be implicated in posttranscriptional regulation of chicken PPARγ. Herein, we focused our attention on the PPARγ1 uORF. Of these five chicken PPARγ transcript variants, PPARγ transcript variant 1 (*PPARγ1*) is highly expressed in various chicken tissues, including abdominal adipose, spleen, and liver (Duan *et al.* 2015), which is consistent with our results showing that PPARγ1 5′ UTR had high reporter activity (Fig. 1A and 1B). Unlike the other two uORF-containing 5′ UTR isoforms, PPARγ1 5′ UTR presented the largest uORF, and its uAUG was in a favorable Kozak consensus context.

To test our speculation, we investigated the effect of PPARγ1 uORF on posttranscriptional regulation of the *Renilla* luciferase reporter gene. We generated a uORF-mutated reporter construct, named PPARγ1-uORF-Mut, by mutating the uAUG to a stop codon UAG (AUG > UAG) of the PPARγ1 uORF by site-directed mutagenesis. Transient transfection and reporter gene assays showed that the luciferase activities of the mutant reporter construct (PPARγ1-uORF-Mut) were 3- and 2.5-fold higher, respectively, than those of the wild-type PPARγ1 5′ UTR reporter construct (PPARγ1-5′UTR-WT) in ICP1 and DF1 cells (*P* < 0.01, Fig. 2B). These results indicate that this uORF functions as an intrinsic repressor for downstream ORF expression. To further understand the molecular mechanism underlying the repressive effect of this uORF, we quantified the relative mRNA levels of *Renilla* luciferase (*hRluc*) in cells transfected with the same amount of PPARγ1-uORF-Mut and PPARγ1-5′UTR-WT, respectively. Surprisingly, in contrast to the reporter gene assay results (Fig. 2B), quantitative real-time RT-PCR showed that transfection of PPARγ1-uORF-Mut resulted in lower *hRluc* mRNA expression compared with the PPARγ1-5′UTR-WT in both ICP1 (*P* < 0.05) and DF1 cells (*P* < 0.01) (Fig. 2C). Thus, the luciferase reporter gene assay and quantitative RT-PCR results together allow us to conclude that *PPARγ1* uORF inhibits *hRluc* translation.

### Inhibition of PPARγ1 translation by the uORF

To exclude the possibility that the effect of this uORF is reporter gene-specific, we generated full-length PPARγ1 expression constructs with either the wild-type or uORF-mutated 5′ UTR (AUG > UAG), termed pcDNA3.1-PPARγ-WT and pcDNA3.1-PPARγ-Mut, respectively. Then, ICP1 cells were transfected with pcDNA3.1-PPARγ-WT or pcDNA3.1-PPARγ-Mut alone, and PPARγ protein expression was assayed by western blot. The western blot analysis showed that PPARγ1 protein levels were significantly higher in the DF1 cells transfected with pcDNA3.1-PPARγ-Mut than with the pcDNA3.1-PPARγ-WT (*P* < 0.01, Fig. 3A and 3B). In parallel, we investigated the *PPARγ1* mRNA expression. Real-time RT-PCR analysis showed that *PPARγ1* mRNA expression levels were significantly lower in both ICP1 and DF1 cells transfected with pcDNA3.1-PPARγ-Mut than with pcDNA3.1-PPARγ-WT (Fig. 3C). These results are consistent with those of the reporter gene assay (Fig. 2C). Collectively, these results indicate that this uORF represses downstream PPARγ1 translation.

**Figure 3.**
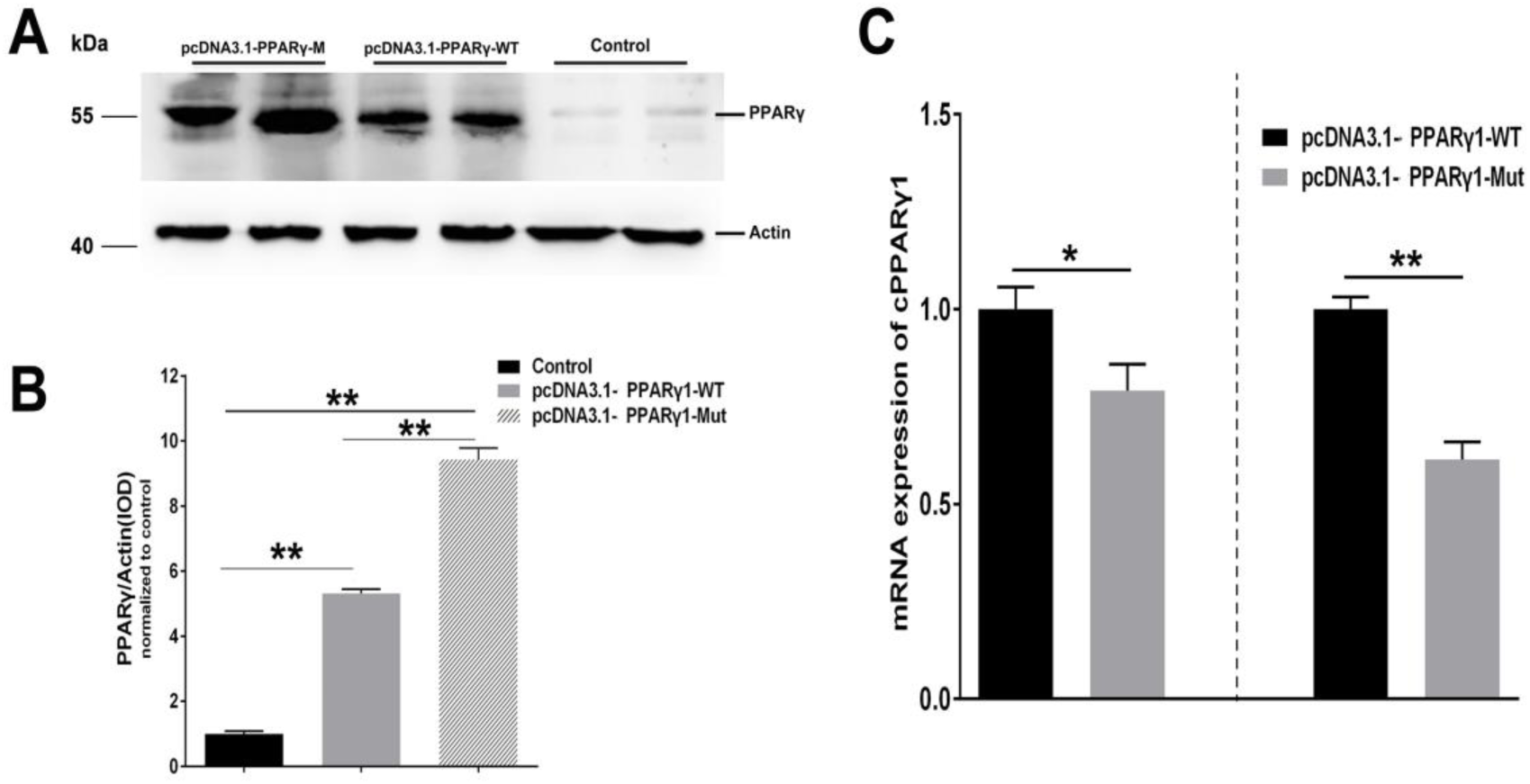
PPARγ1 translation is inhibited by its 5′ UTR uORF. (A) Detection of PPARγ1 protein levels. Equal amounts of the total cell lysates from the ICP1 cells transfected with either pcDNA3.1-PPARγ-WT or pcDNA3.1-PPARγ-Mut were separated and immunoblotted with an anti-PPARγ antibody. Actin was used as a loading control. (B) Quantification of PPARγ1 protein expression. Band intensities were measured by ImageJ software normalized to actin loading control. Data represent Mean ± SEM. PPARγ1 protein expression was higher in the cells transfected with pcDNA3.1-PPARγ-Mut than in the cells transfected with the pcDNA3.1-PPARγ-WT (** P < 0.01, Student’s t-test). (C) Quantification of PPARγ1 mRNA by real-time RT-PCR in the ICP1s and DF1 cells transfected with the indicated constructs. PPARγ1 mRNA levels were normalized to the expression of the cells transfected with pcDNA3.1-PPARγ-WT. Data were expressed as the mean ± SEM, NONO was used as the internal mRNA control. n ≥ 3, *P < 0.05; ** P < 0.01, Student’s t-test.

### No effect of PPARγ1 uORF on mRNA stability

Our results showed that the uORF mutation resulted in reduced mRNA expression levels of *hRluc* and *PPARγ1* (Fig. 2C and 3C). There are two possibilities to explain this. First, the uORF mutation may affect mRNA stability. Previous studies have indicated that uORF can reduce mRNA expression via mRNA destabilization (Dikstein 2012; Dvir *et al.* 2013). The other possibility is that the cloned chicken PPARγ1 5′ UTR in our 5′ UTR reporters and PPARγ expression vectors may contain promoter activity, and uORF mutation may lead to reduced promoter activity. To test whether this uORF mutation affected mRNA stability, using real-time RT-PCR, we determined the mRNA decay rate of *hRluc* in cells transfected with PPARγ1-5′UTR-WT or PPARγ1-5′UTR-Mut at 0, 3, 6, 9, and 12 h following treatment with actinomycin D. As shown in Fig. 4, no significant difference in *hRluc* mRNA half-life was observed over a 12-h period between cells transfected with PPARγ1-5′UTR-WT or PPARγ1-5′UTR-Mut (PPARγ1-5′UTR-WT: 6.43 h; PPARγ1-5′UTR-Mut: 5.90 h (*P* = 0.4146). These results indicate that this uORF had no obvious effect on mRNA stability.

**Figure 4.**
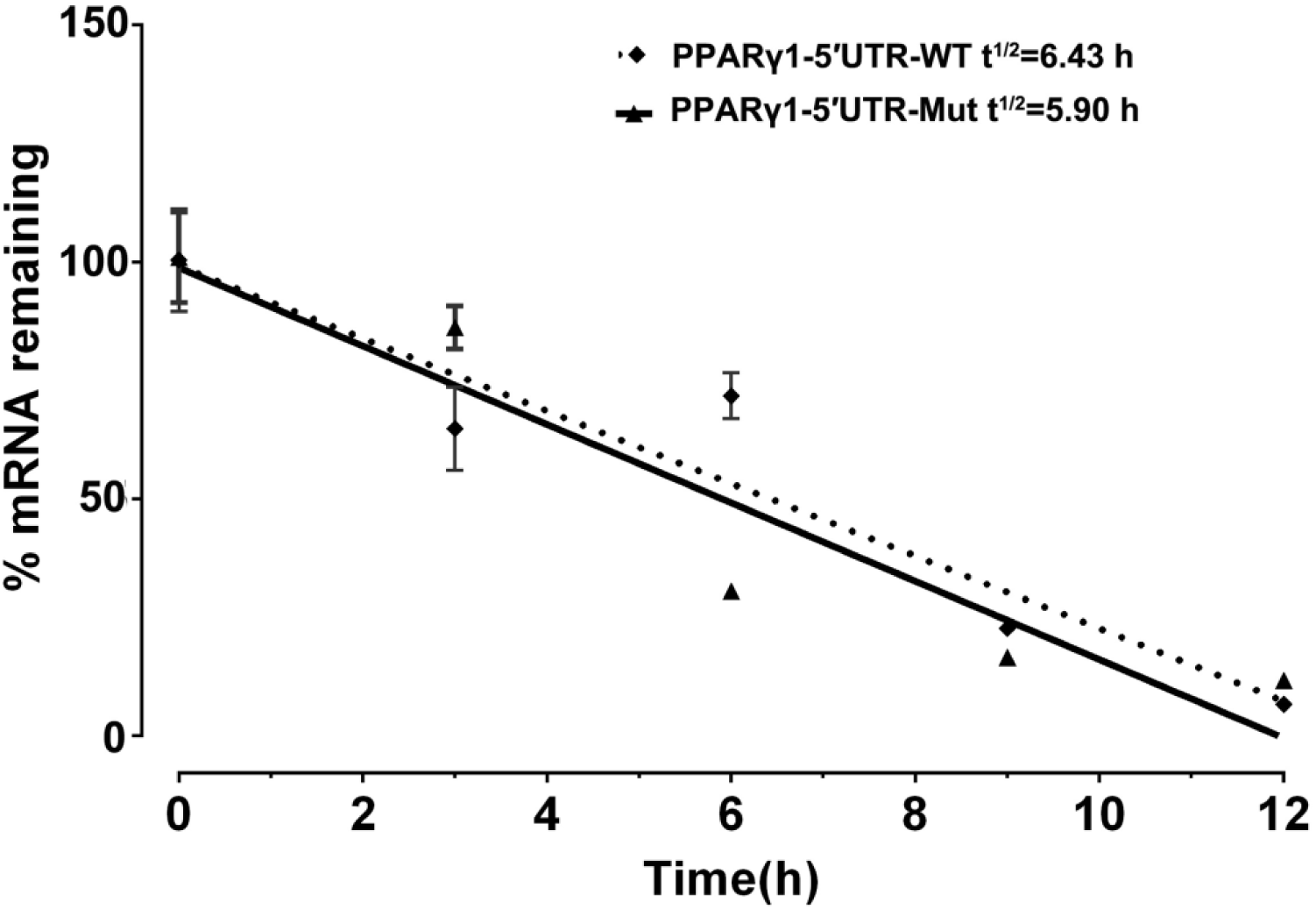
Effect of uORF mutation on hRluc mRNA stability. ICP1 cells were transiently transfected with PPARγ1-5′UTR-WT or PPARγ1-5′UTR-Mut, 48 h post-transfection, hRluc mRNA remaining after a 12 h time-course treatment with Actinomycin D was measured by real-time RT-PCR and calculated as a percentage of the level measured at time zero (0 h). Linear regression analysis was used to determine the half-life of the hRluc mRNA (t1/2), the time required for degrading 50% of the existing hRluc mRNA molecules at 0 h. No differences in relative mRNA decay rate were observed between the cells transfected with PPARγ1-5′UTR-WT and PPARγ1-5′UTR-Mut. Data are expressed as the mean ± SEM relative to NONO expression.

### The effect of uORF mutation on promoter activity

The genomic region corresponding to the 5′ UTR is usually part of the promoter. Our previous study demonstrated that the 108-bp sequence downstream of the transcription start site of *PPARγ1*, which is part of the 5′ UTR, had the highest promoter activity (Cui *et al.* 2018). To test whether the cloned PPARγ1 5′ UTR had promoter activity and whether the uORF mutation reduced it, we cloned DNA sequences corresponding to wild-type and uORF-mutated 5′ UTRs of PPARγ1 into luciferase reporter vector pGL3-Basic, named pGL3-PPARγ1-WT and pGL3-PPARγ1-Mut, respectively. A reporter gene assay showed that the pGL3-PPARγ1-WT and pGL3-PPARγ1-Mut displayed 111- and 90-fold higher luciferase reporter activity, respectively, than the pGL3-Basic empty vector in DF1 cells, and 180- and 120-fold higher luciferase reporter activity, respectively, than the pGL3-Basic empty vector in ICP1 cells, suggesting that the cloned PPARγ1 5′ UTR has promoter activity. By comparison, pGL3-PPARγ1-Mut showed significantly lower luciferase activity than pGL3-PPARγ1-WT in DF1 cells (Fig. 5A, *P* < 0.05) and ICP1 cells (Fig. 5B, *P* < 0.01). These results demonstrated that the cloned PPARγ1 5′ UTR had strong promoter activity and that the uORF mutation can result in reduced promoter activity. These findings explain why the mRNA expression levels of *hRluc* and *PPARγ1* were reduced in the above study (Fig. 2C and 3C).

**Figure 5.**
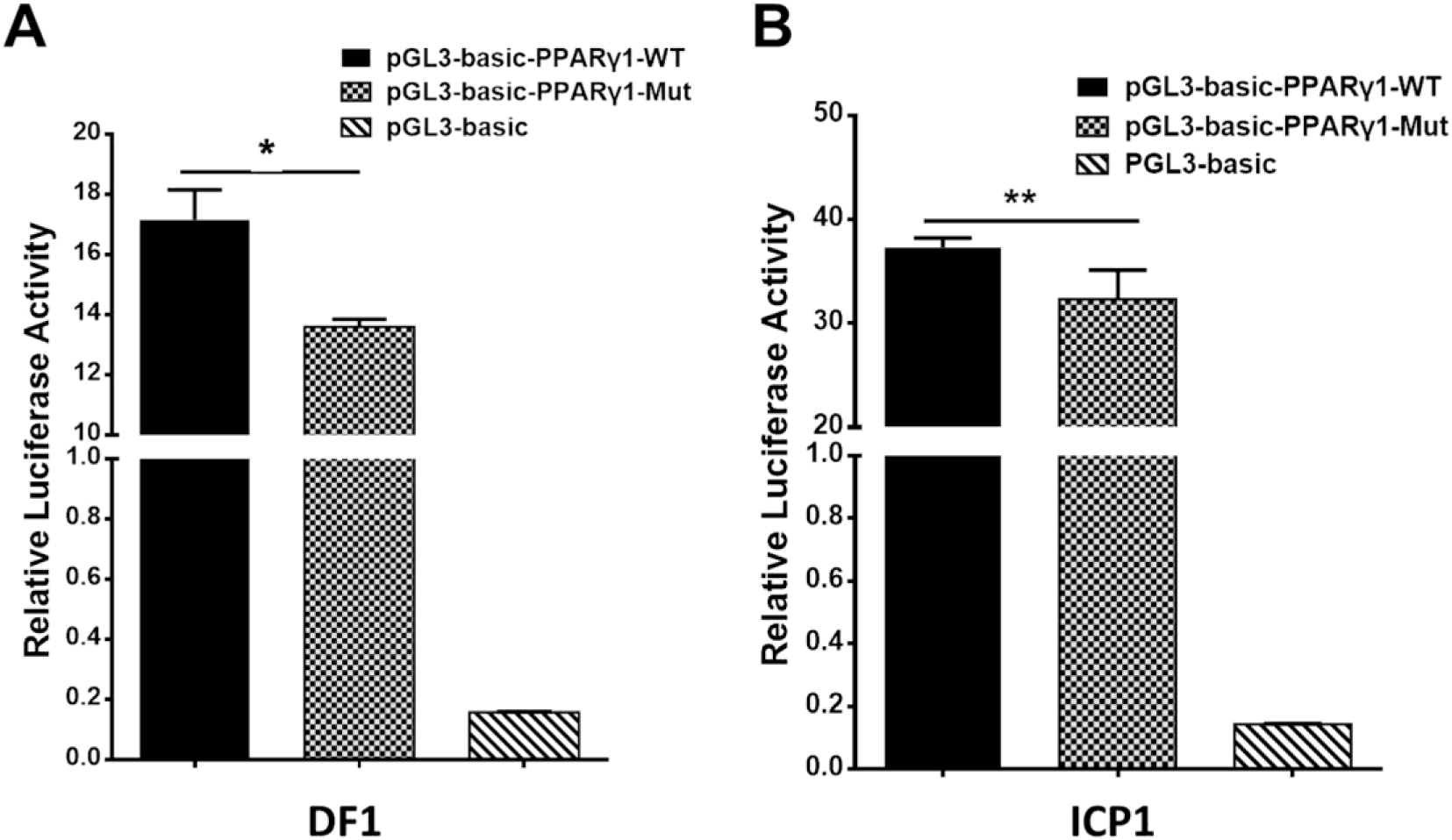
The promoter activity analysis of the DNA sequences corresponding wild-type and uORF-mutated 5′ UTRs of PPARγ1. The DNA sequences corresponding wild-type and uORF-mutated 5′ UTRs of PPARγ1 were cloned into luciferase reporter vector pGL3-basic to yield pGL3-PPARγ1-WT and pGL3-PPARγ1-Mut, respectively. The indicated reporters along with the pRL-TK Renilla luciferase vector were transiently transfected into DF1 (A) and ICP1 cells (B), and the luciferase activity was determined at 48 h after transfection. The pRL-TK vector was used for normalization of transfection efficiency. All data represent the mean ± SEM. ** p < 0.01, Student’s t-test.

### The effect of the uORF on *in vitro* translation of PPARγ1

The results reported here suggest that at the mRNA level, the uORF represses PPARγ1 translation (Fig. 3A and 3B). To further validate this finding, we performed an *in vitro* transcription and translation assay. For *in vitro* transcription, equal amounts (1 μg) of linearized pcDNA3.1-PPARγ1-WT or pcDNA3.1-PPARγ1-Mut were used as templates to produce the *PPARγ1* mRNA with wild-type and uORF-mutated 5′ UTR using the T7 RiboMAX Large Scale RNA Production System. The results showed that, as expected, pcDNA3.1-PPARγ1-WT and pcDNA3.1-PPARγ1-Mut produced almost the same amount of PPARγ1 mRNA (Fig. 6A). Equal amounts of the transcribed PPARγ1 mRNA produced from pcDNA3.1-PPARγ1-WT and pcDNA3.1-PPARγ1-Mut were used for the *in vitro* translation assay. The *in vitro* translation assay results showed that more PPARγ1 protein was synthesized with PPARγ1 mRNA from pcDNA3.1-PPARγ1-Mut than from pcDNA3.1-PPARγ1-WT (Fig. 6B). Together, these results indicate that the uORF represses PPARγ1 translation.

**Figure 6.**
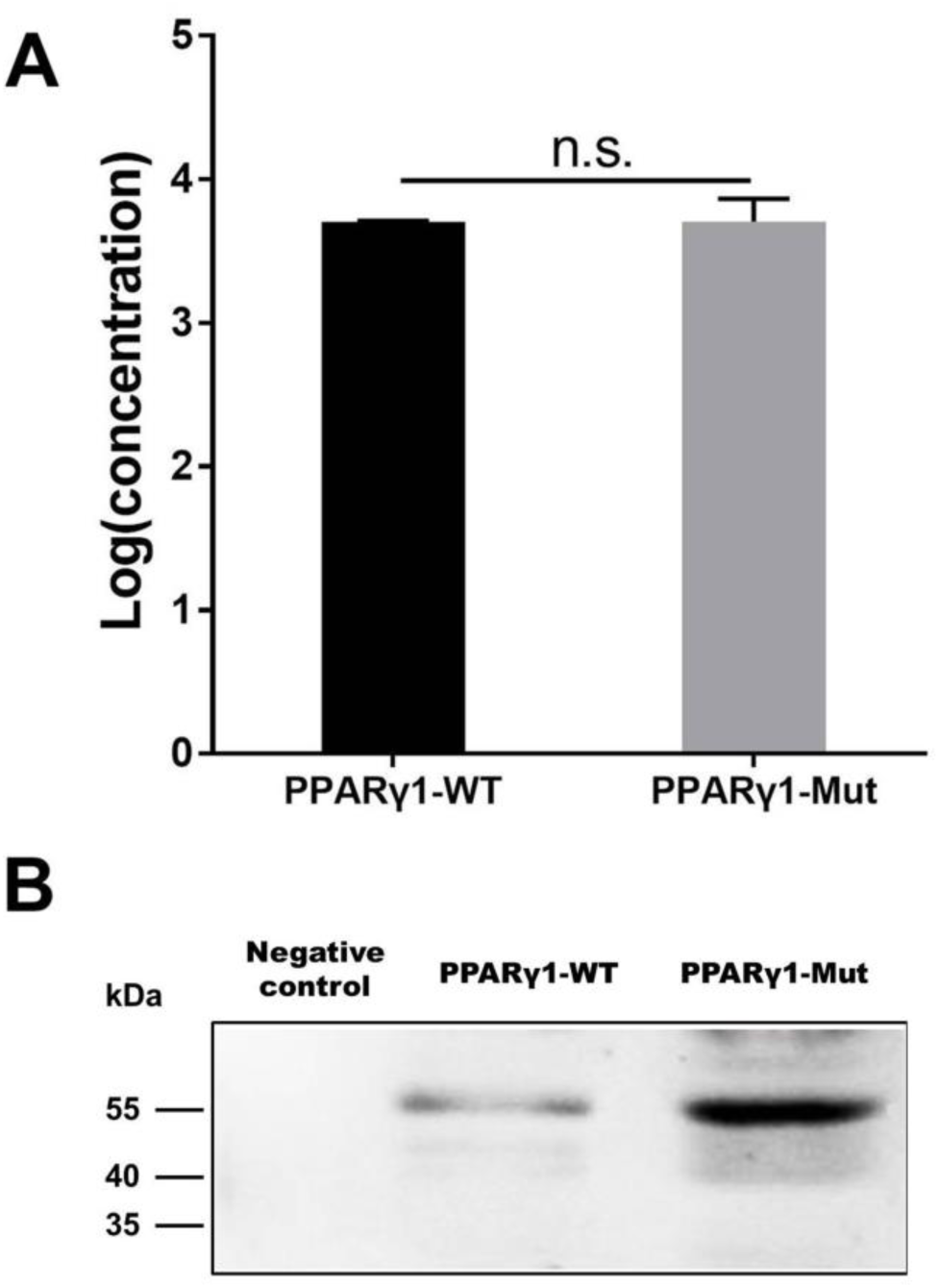
The uORF represses in vitro PPARγ1 translation. (A) In vitro transcribed PPARγ1 mRNAs from the wild-type and uORF-mutant PPARγ1 expression vectors (pcDNA3.1-PPARγ1-WT and pcDNA3.1-PPARγ1-Mut) were analyzed by quantitative real-time RT-PCR. No difference in PPARγ1 mRNA was observed. Data were expressed as the mean ± SEM, n.s., not significant, Student’s t-test. (B) Equal amounts of the in vitro transcribed mRNAs (2 μg) were used for in vitro translation. Note that the uORF strongly represses PPARγ1 translation. A in vitro translation reaction without RNA template was used as a negative control.

### The PPARγ1 uORF can be translated

To gain insight into the molecular mechanism by which the uORF represses translation, we tested whether the uAUG of PPARγ1 uORF was used for translation initiation. We generated a construct in which the uORF was fused in frame with the enhanced green fluorescent protein (EGFP) coding sequences, with no intervening in-frame stop codons, and named it pcDNA3.1-uORF-EGFP; pcDNA3.1-EGFP was used as a positive control. The ICP1 cells were transiently transfected with pcDNA3.1-uORF-EGFP or pcDNA3.1-EGFP and examined by microscopy and western blotting. Microscopy showed that the cells transfected with either pcDNA3.1-uORF-EGFP or pcDNA3.1-EGFP displayed GFP fluorescence (Fig. 7A). Comparatively, GFP fluorescence intensity was lower in the cells transfected with pcDNA3.1-uORF-EGFP than with pcDNA3.1-EGFP (Fig. 7A). Consistent with these findings, western blot analysis showed that the uORF-EGFP fusion protein was expressed but at a lower level in the cells transfected with pcDNA3.1-uORF-EGFP compared with that in the cells transfected with pcDNA3.1-EGFP (Fig. 7B). Collectively, these data suggest that translation can indeed be initiated at the uAUG.

**Figure 7.**
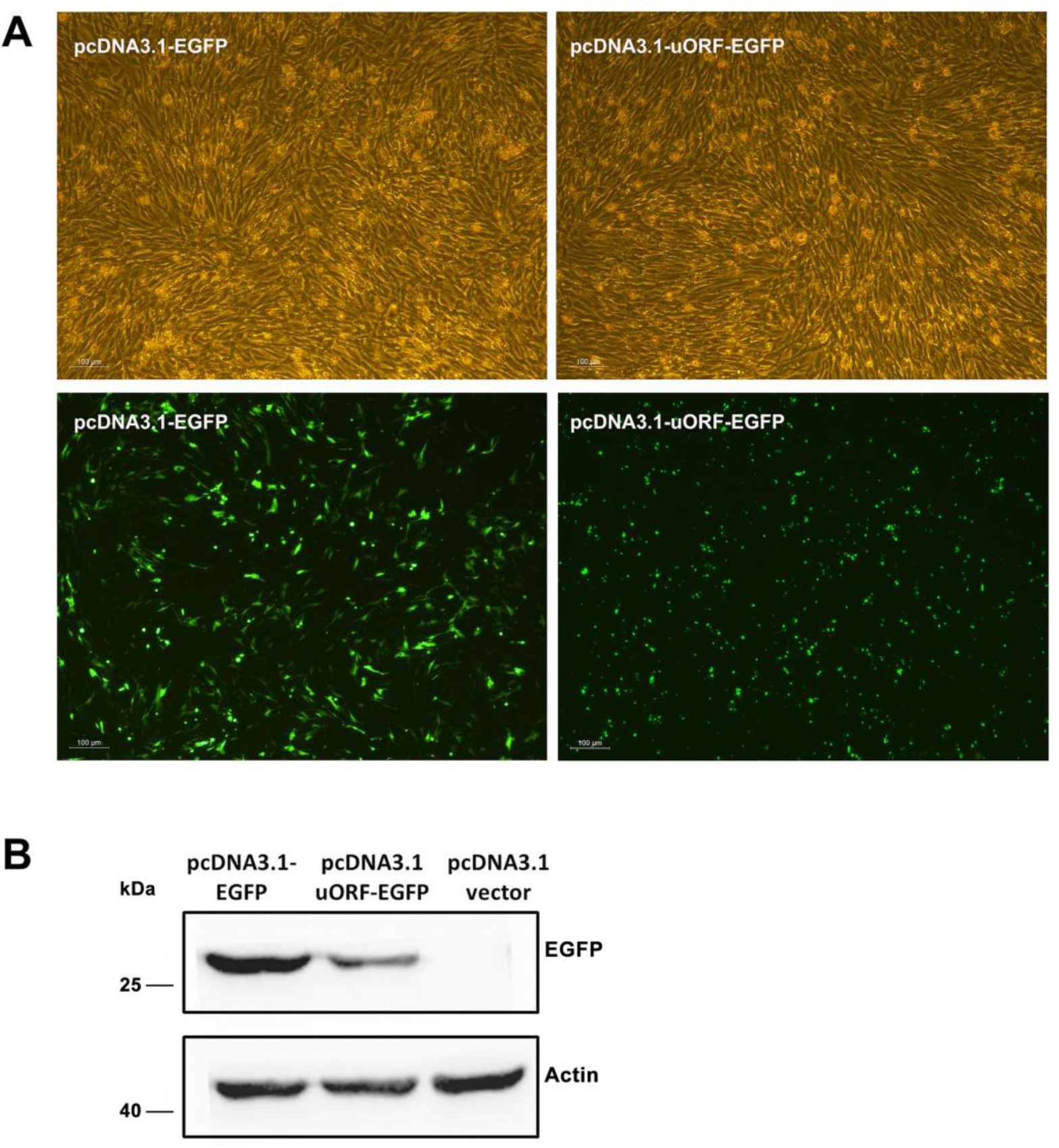
Translation can be initiated at the uAUG of the PPARγ1 uORF. (A) The pcDNA3.1-EGFP and pcDNA3.1-uORF-EGFP were respectively transiently transfected into ICP1 cells, 48 h post-transfection, the green fluorescence signal was visualized under a fluorescence microscope. (B) Lysates from the cells transfected with pcDNA3.1-EGFP and pcDNA3.1-uORF-EGFP, EGFP or uORF-EGFP fusion protein was immunoblotted with an anti-EGFP antibody. Actin was used as a loading control.

## Discussion

Investigating the molecular mechanisms that control PPARγ expression is critical for understanding adipogenesis, as well as pathological conditions such as obesity and diabetes. In the present study, we investigated PPARγ posttranscriptional regulation by 5′ UTR. We demonstrated that a uORF, which is absent in human and mouse PPARγ transcripts, represses chicken PPARγ transcript variant 1 translation. To our knowledge, this is the first report of a uORF regulating *PPARγ* gene expression.

In the present study, we demonstrated that the five chicken PPARγ 5′ UTR isoforms exerted different effects on the reporter gene activity (Fig. 1A and 1B), and further study showed the 5′ UTR uORF of PPARγ1 represses reporter gene and PPARγ1 translation. Sequence analysis revealed that the uAUG of the PPARγ1 uORF resides in a favorable Kozak sequence context, which is the most efficient context for ribosome recognition and initiation of translation (Fig. 2A). In agreement with the bioinformatics prediction, we demonstrated that the uAUG could serve as a translation start site (Fig. 7A and 7B). Furthermore, secondary structural analysis showed there was a stable loop structure within the uORF (Fig. S1), and the uORF mutation (AUG > UAG) was not able to alter the secondary structure of PPARγ1 5′ UTR (Fig. S1). Thus, we could rule out an effect of secondary structure alteration on PPARγ1 expression.

The 17-amino acid PPARγ1 uORF peptide was analyzed using the PSIPRED protein sequence analysis workbench. It was predicted to be a disordered peptide (Romero *et al.* 2001; Buchan *et al.* 2013). Disordered peptides are enriched with residues Gly, Pro, Arg, and Ser, which are potential targets for phosphorylation that could promote ribosome stalling during translation elongation or termination (Hayden and Jorgensen 2007; Johansson *et al.* 2011; Koutmou *et al.* 2015), which may explain why EGFP expression was lower in the cells transfected with pcDNA3.1-uORF-EGFP than with pcDNA3.1-EGFP (Fig. 7A and 7B).

An increasing number of uORF-encoded peptides have been identified and shown to repress the downstream ORF expression by triggering ribosome stalling or suppressing reinitiation (Wilson and Beckmann 2011; Ito and Chiba 2013; Starck *et al.* 2016). Our data demonstrated that the PPARγ1 uORF repressed downstream ORF expression and that it could be translated. This raised the question of whether this uORF-encoded peptide represses downstream ORF translation. To this end, we constructed a uORF expression vector, pcDNA3.1-uORF, and co-transfected pcDNA3.1-uORF and PPARγ1-5′UTR-WT or PPARγ1-5′UTR-Mut into DF1 cells. Unexpectedly, reporter gene assays showed that transfection of pcDNA3.1-uORF increased reporter gene activities of both PPARγ1-5′UTR-WT and PPARγ1-5′UTR-Mut (Fig. S2). This result suggested that this uORF-encoded peptide may repress the downstream PPARγ1 translation in *cis*, but not in *trans.* It has been reported that several uORF-encoded peptides act in *cis* on the ribosome during their own translation to stall translation. Arrest of translation can occur either during translation elongation, as seen for SecM (Tsai *et al.* 2014) and VemP (Vazquez-Laslop *et al.* 2008), or during translation termination; for example, in the tryptophanase C (TnaC) (Gong *et al.* 2001) and S-adenosyl-methionine decarboxylase (SAM-DC) (Raney *et al.* 2002).

Based on our data, we speculated that uORF repressed PPARγ1 translation by two possible mechanisms. The first was ribosome stalling (Fig. 8A), in which uATG is recognized by the scanning 40*S* ribosomal subunit and associated initiation factors, the uORF is translated, and the nascent peptide stalls the ribosome in the ribosome exit tunnel, thereby hampering the progression of upstream ribosomes (Wilson 2011; Brandman *et al.* 2012; Wilson *et al.* 2016). Only a tiny minority of ribosomes may leaky-scan the uORF start codon and translate the PPARγ1 coding sequence. Consequently, the translational efficiency of PPARγ1 is dramatically attenuated.

**Figure 8.**
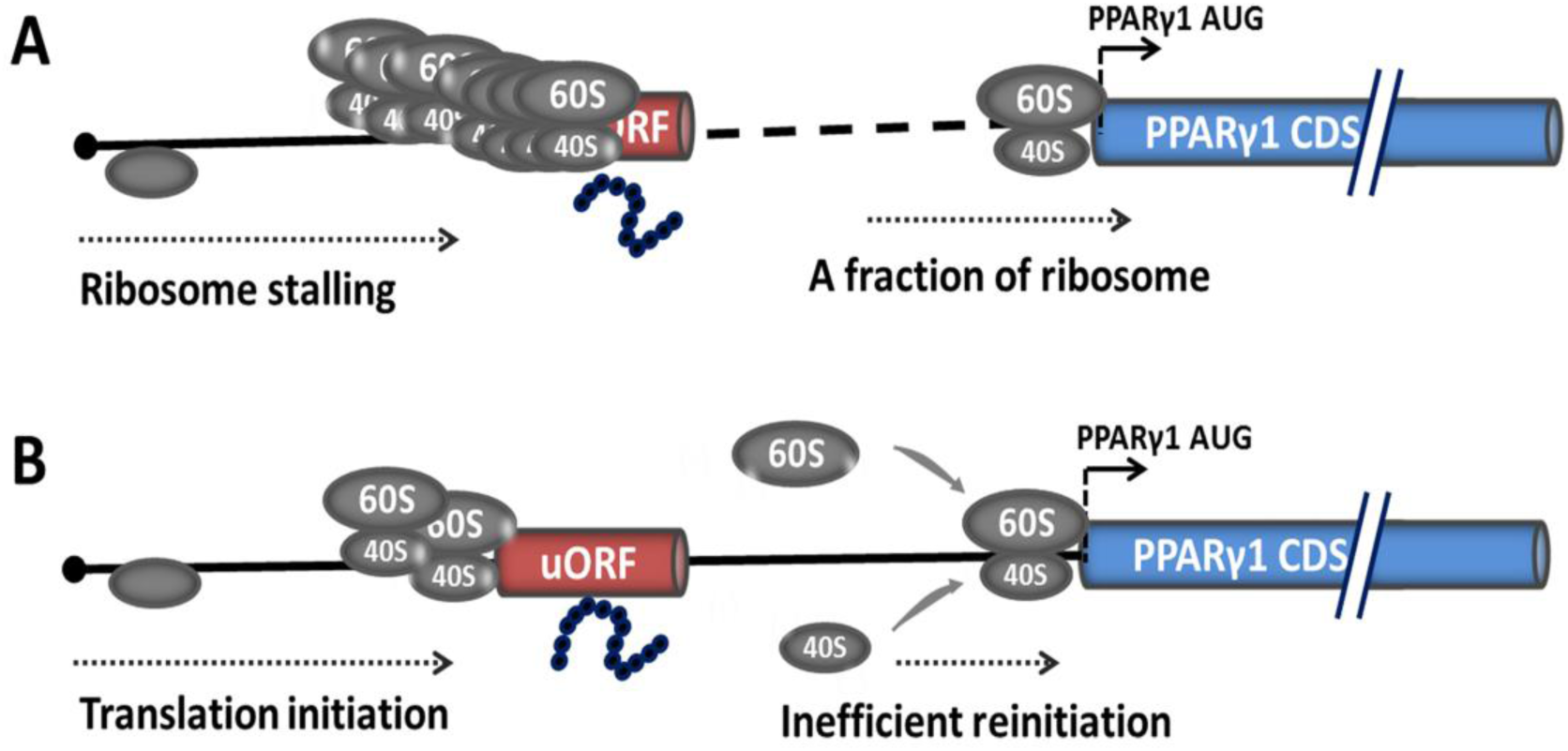
Potential models for uORF-mediated PPARγ1 translational inhibition. Translational inhibition of PPARγ1 may be due to uORF-mediated ribosome stalling (A) or inefficient reinitiation at the authentic start codon of PPARγ1 (B).

The other possible mechanism is translational reinitiation, in which ribosomes translate the uORF and remain associated with the mRNA, continue scanning, and reinitiate further downstream at either a proximal or distal AUG codon (Fig. 8B). However, reinitiation efficiency is substantially reduced (Roy *et al.* 2010; Hinnebusch *et al.* 2016) and translation of PPARγ1 inhibited. Recently, some nascent peptides of uORFs have been reported to be involved in the suppression of reinitiation (Ito and Chiba 2013; Seefeldt *et al.* 2015). We speculate that the uORF-encoded peptide may contribute to suppression of PPARγ1 translation.

In addition, several studies have implicated that uORF-containing mRNA has the potential to trigger the nonsense-mediated decay (NMD) pathway. NMD is one of the better characterized posttranscriptional control mechanisms, whereby transcripts harboring premature translation termination codons are degraded (Mendell *et al.* 2004). In the present study, we detected no significant effect of uORF mutation on *hRluc* mRNA stability (Fig. 4 and Fig. 6A). Therefore, we can rule out the possibility that PPARγ1 uORF modulates PPARγ1 expression by triggering the NMD pathway.

PPARγ is a master regulator of adipogenesis, whole-body lipid metabolism, and insulin sensitivity. Accumulating evidence shows that adipogenesis and lipid metabolism are different between mammals and chickens (Prigge and Grande 1971; Ji *et al.* 2012). For example, unlike that in mammals, chicken adipocyte lipolysis is almost exclusively regulated by glucagon, and chicken adipose tissue is not sensitive to insulin (Dupont *et al.* 2012; Ji *et al.* 2012; Wang *et al.* 2017). In the present study, we demonstrated that the uORF represses chicken PPARγ1 translation, and bioinformatics analysis showed that PPARγ 5′ UTRs have very low sequence similarity between humans, mice, and chickens; no uORF element is present in the 5′ UTRs of human and mouse PPARγ transcripts. Our data suggest that the posttranscriptional regulation of the *PPARγ* gene by the 5′ UTR differs between mammals and chickens, which may contribute to differences in adipogenesis, adipose development, and insulin sensitivity between mammals and chickens. It is worth further exploring the roles and underlying mechanisms of PPARγ 5′ UTRs. A better understanding of PPARγ 5′ UTRs may provide clues for treating obesity, type 2 diabetes, and insulin resistance.

In summary, for the first time, we demonstrated that a uORF represses chicken PPARγ1 translation.

## Acknowledgments

We thank Louise Adam, ELS(D), from Liwen Bianji, Edanz Editing China (www.liwenbianji.cn/ac) for editing the English text of a draft of this manuscript. This work was supported by the National Natural Science Foundation of China (No. 31572392) and the China Agriculture Research System (No. CARS-41).

## Conflict of interest

The authors declare that they have no conflicts of interest with the contents of this article.

## Author contributions

Y Chu designed the experiments, collected and analyzed data, and wrote the manuscript. J Huang, G Ma, T Cui, X Yan and H Li participated in scientific discussions and provided technical assistances. N Wang supervised the study and wrote the manuscript with Y Chu.

## Supplementary Materials

**Supplementary figure 1.**
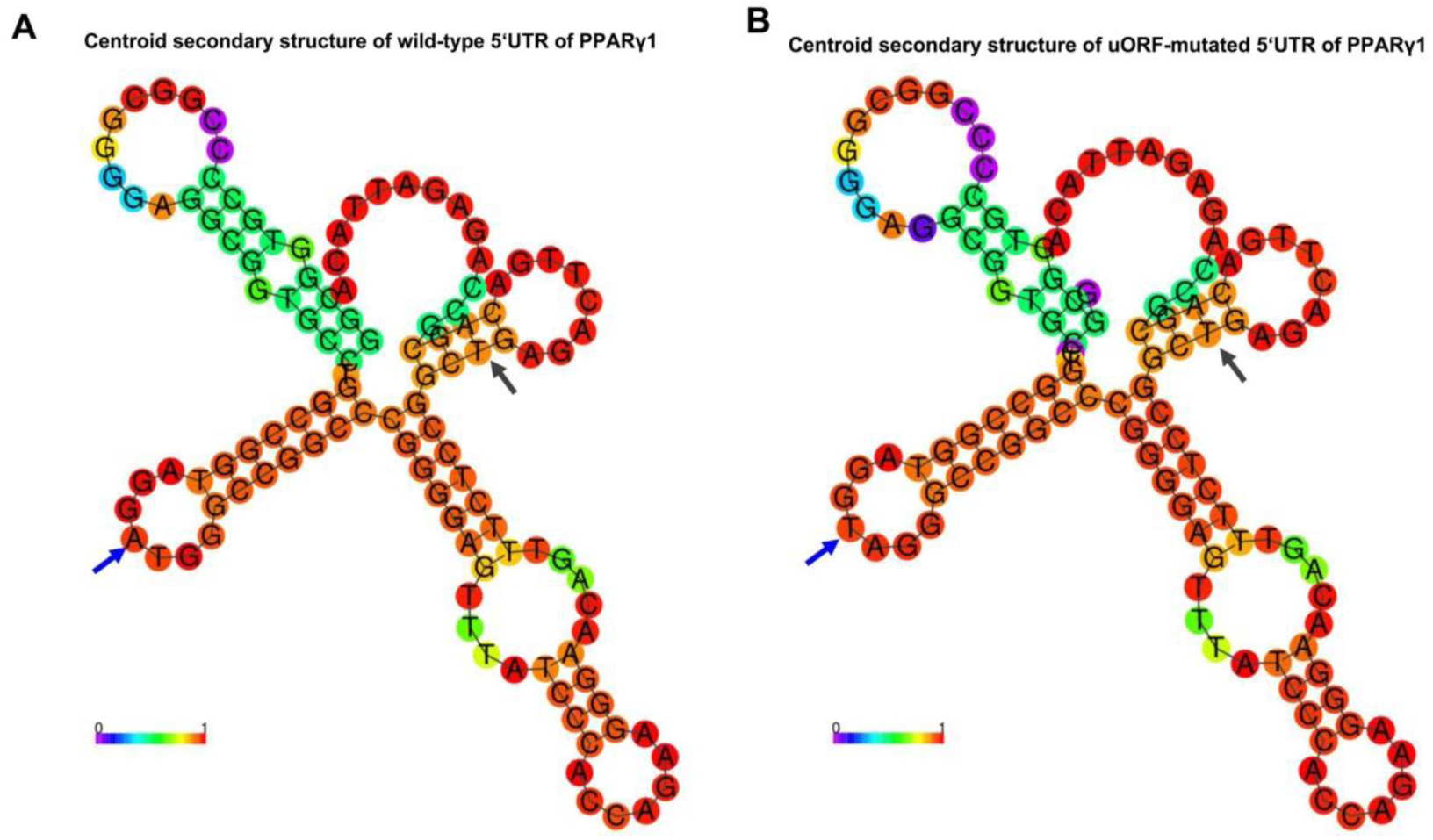
Predicted secondary structure of the wild-type and uORF-mutant 5′ UTR of chicken *PPARγ1* mRNA. RNAfold 2.0 (http://rna.tbi.univie.ac.at/cgi-bin/RNAfold.cgi) was used for structure prediction. The centroid structures encoding base pair probabilities are shown. The bases are colored in violet (0) for low and in *red* (1) for high base-pairing probabilities. The uORF within the 5′UTR are indicated by blue (5′ terminal) and black (3′ terminal) *arrows*.

**Supplementary figure 2.**
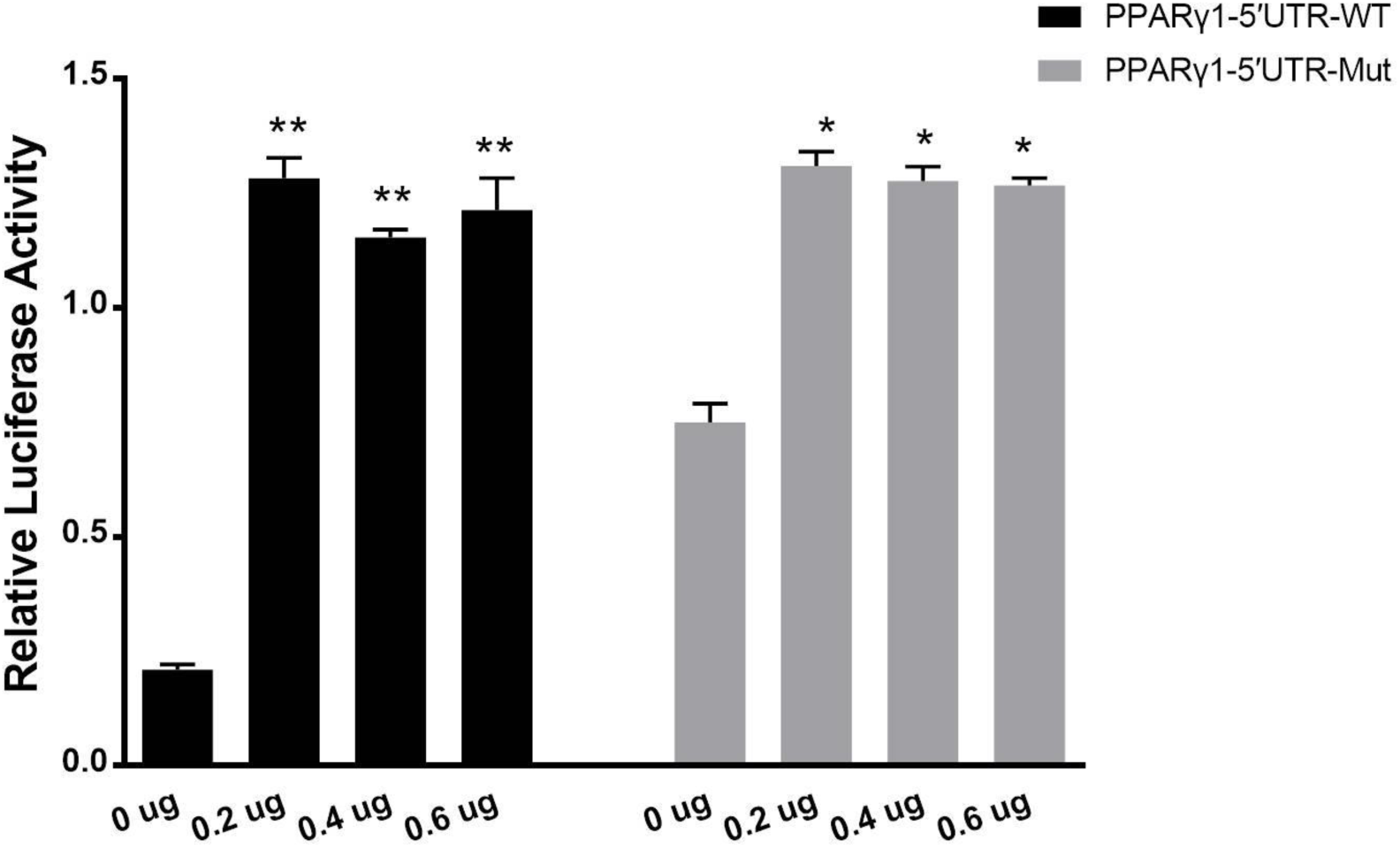
The uORF-encoded peptide does not repress the downstream reporter gene translation in *trans*. The indicated amounts of the uORF expression vector (pcDNA3.1-uORF) and either PPARγ1-5′UTR-WT or PPARγ1-5′UTR-Mut were contransfected into DF1 cells, respectively, and the luciferase activity was determined at 48 h after transfection. All data represent the mean ± SEM. * *p* < 0.05, Student’s *t*-test.

